# Immunity can impose a reproduction-survival tradeoff on human malaria parasites

**DOI:** 10.1101/2025.01.27.635035

**Authors:** Denis D. Patterson, Lauren M. Childs, Isaac J. Stopard, Nakul Chitnis, Sergio Serrato-Arroyo, Megan Greischar

## Abstract

Many pathogenic organisms produce specialized life stages for within-host multiplication versus onward transmission, including malaria parasites. Traits that enable faster multiplication—including limited investment into transmission stage production—should put host health at greater risk (all else equal). Yet it is not clear why parasites do not evolve ever faster multiplication rates, since malaria parasites do not appear to adhere to tradeoffs between the rate and duration of transmission that are classically predicted to constrain parasite evolution. To address this puzzle, we introduce an age-of-infection structured within-host mathematical model incorporating dynamic immune clearance to investigate potential tradeoffs and understand how parasites optimize their transmission investment. When investment is constant across all ages of infection, increased transmission investment reduces infection duration and parasite fitness, with optimal investment occurring at a relatively low value (around 5%), far lower than the optimum recovered from models that lack dynamic feedbacks between parasite investment and immune clearance. For age-varying strategies, our model shows that malaria parasites can enhance their fitness by delaying transmission investment to allow for faster within-host multiplication initially. Our results indicate that adaptive immunity can impose a survival-reproduction tradeoff that explains why malaria parasites cannot evolve ever faster within-host multiplication. Our theoretical framework provides a basis for understanding how transmission investment strategies alter the timing of infectiousness over the lifespan of malaria infections, with implications for parasite evolution in response to control efforts.

## 1 Introduction

Classic theory predicts that tradeoffs between the rate and duration of transmission constrain parasite evolution towards ever faster multiplication rates within the host Anderson and May (1982); Frank (1996). The idea of such transmission-duration tradeoffs has found support in HIV infections Fraser et al. (2014); Payne et al. (2014), where higher viral loads increase the odds of transmission while hastening the end of infection via progression to AIDS. Tradeoffs between the rate and duration of transmission could arise from two distinct but similar mechanisms: transmission-virulence or transmission-recovery tradeoffs. The assumption of a transmission-virulence tradeoff, where infection-induced host mortality (virulence) is thought to limit transmission duration, underpins important theory, e.g., on pathogen evolution in response to vaccines Gandon et al. (2001); Miller and Metcalf (2022). However, the generality of transmission-virulence tradeoffs is debated due in part to the low case fatality rates of many human pathogens Bull and Lauring (2014). Similarly, faster within-host multiplication can enhance transmission but lead to faster host recovery, imposing a transmission-recovery tradeoff. For example, larger viral loads in human dengue infections increase rates of transmission to mosquitoes Nguyen et al. (2013) but hastening clearance by the immune system Ben-Shachar and Koelle (2018).

Such tradeoffs cannot always explain what prevents the evolution of faster multiplication rates of multiplication, since some pathogenic organisms, including human malaria parasites (*Plasmodium falciparum*), do not appear subject to transmission-virulence or transmission-recovery tradeoffs. Despite their large health burden, most human malaria infections are subclinical, particularly in high transmission regions where frequent exposure speeds development of immunity that reduces the symptoms of disease Lindblade et al. (2013). Within human hosts, parasite abundance can increase substantially without significantly increasing the risk of severe infection outcomes Cunnington et al. (2013). Faster parasite multiplication extends rather than abbreviates the time until immune clearance Mackinnon and Read (2004), which models suggest results from complicated interactions between parasites’ antigenic repertoire and cross-reactive immune responses Klein et al. (2014); Childs and Buckee (2015). Thus, classic transmission-duration tradeoffs cannot explain what prevents malaria parasites from evolving faster multiplication rates.

Life history theory (LHT) offers a distinct prediction from the transmission-duration tradeoff for what prevents organisms from evolving ever greater reproduction rates: a tradeoff between investment into reproduction and survival. Sometimes, the predictions overlap, as for many pathogenic organisms, the LHT prediction is analogous to the transmission-duration tradeoff, where seeding new infections (transmission) and persistence within the host represent reproduction and survival, respectively. However, the predictions emerging from LHT are not always equivalent to those following from a transmission-duration tradeoff (Figure 1). Some pathogenic organisms, like malaria parasites, produce specialized forms for multiplication within the host versus onward transmission. For such organisms, investment into transmission (reproduction) comes at the cost of multiplication within the host, but the constraints on the evolution of transmission investment remain unclear. Under a transmission-duration tradeoff, greater investment into transmission (i.e., the production of specialized transmission stages) should slow within-host multiplication and extend infection duration. In contrast, LHT predicts that greater investment into transmission will limit survival, reducing infection length. Incorporated into multiscale models, these divergent predictions yield qualitatively different outcomes for the evolution of transmission investment in response to control efforts, with consequences for the severity of individual infections and for epidemic spread Greischar et al. (2019).

**Figure 1:**
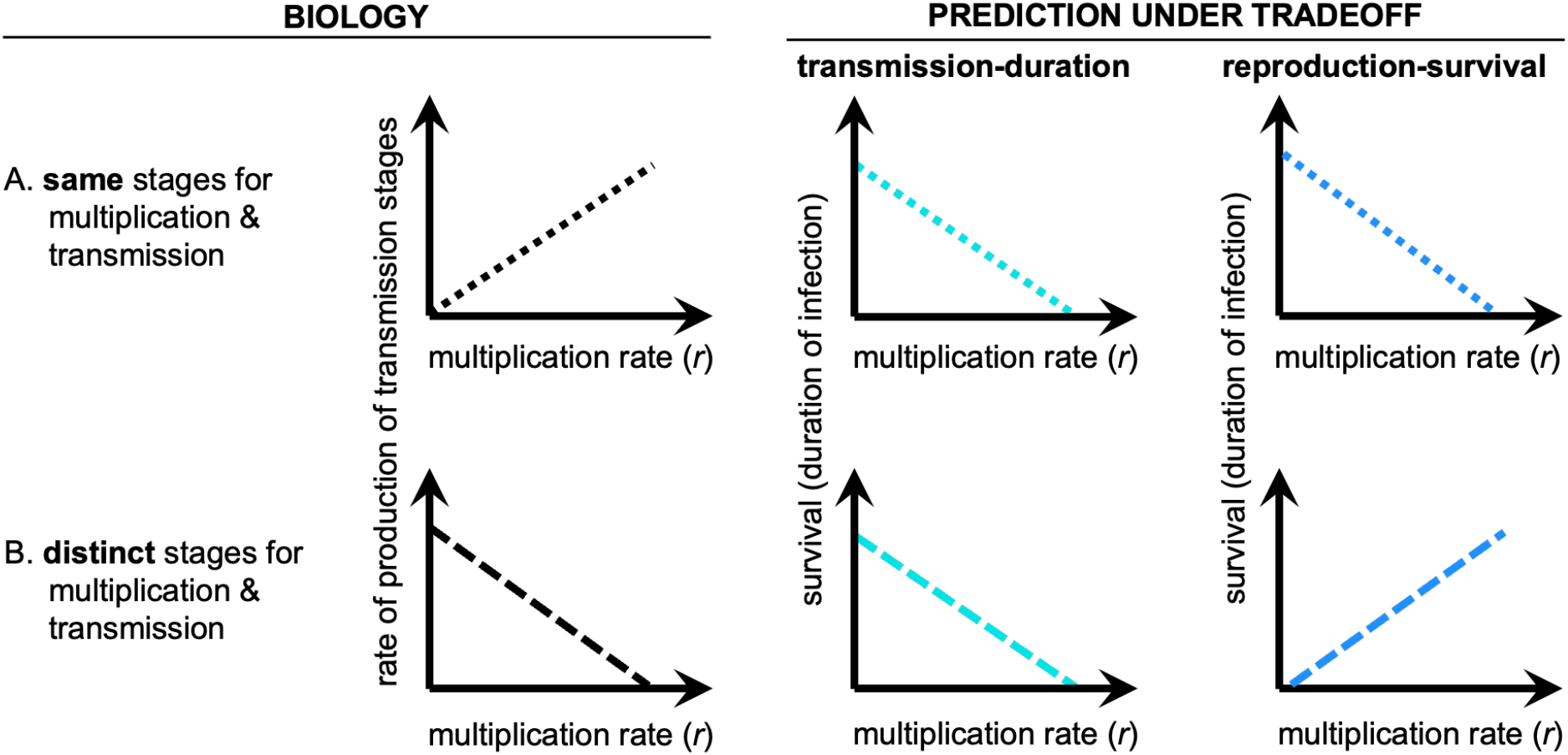
Hypothesized transmission-duration tradeoffs and reproduction-survival tradeoffs may generate identical or opposing predictions depending on the biology of the pathogenic organism (left panels). The transmission-duration tradeoff (middle panels) assumes that transmission (i.e., reproduction) is enhanced by a trait such as multiplication rate that also shortens infection by hastening host recovery or mortality. The reproduction-survival tradeoff (right panels) assumes that investing in reproduction (e.g., producing transmission stages) comes at the cost of continued survival. (A) When the same stages accomplish multiplication and transmission (dotted lines), the transmission-duration and reproduction-survival tradeoffs yield identical predictions that continued persistence within the host (survival) declines with increasing multiplication rate. (B) When parasites produce specialized transmission stages (dashed lines), that production necessarily reduces replication rates, generating opposing predictions for how survival within the host scales with multiplication rate. Survival should either decline with increasing multiplication rates (transmission-duration tradeoff) or increase with multiplication rates due to decreasing production of transmission stages (reproduction-survival tradeoff).

Distinguishing between the competing predictions of LHT and the transmission-duration tradeoff remains an open question in many systems, including malaria. Working with human infection data is fraught with challenges, such as inferring correct patterns of transmission investment. Methods for inference often fail spectacularly when applied to simulated infection data Greischar et al. (2016b), and existing methods for determining transmission investment from human malaria infection data Diebner et al. (2000); Eichner et al. (2001) have not yet been validated against simulated data. Despite the methodological challenges, multiple lines of evidence suggest that transmission investment changes over the course of infection Diebner et al. (2000); Eichner et al. (2001); Greischar et al. (2016b). The advantages of increasing transmission investment are thought to vary as conditions deteriorate within the host Pollitt et al. (2011); Greischar et al. (2016a), and the survival consequences of investment should likewise vary. Yet, linking features of time-varying patterns of transmission investment with survival outcomes is a nontrivial statistical problem. To circumvent these challenges, we extend an existing model of within-host infection dynamics for human malaria parasites (*Plasmodium falciparum*) to include realistic immune clearance over the course of the infection and identify what tradeoffs emerge. Consistent with LHT predictions, we find that increased transmission investment typically reduces infection duration and parasite fitness, leading to a much lower optimal transmission investment than previously predicted Greischar et al. (2019). While increased transmission investment slows the increase in immune removal, it also comes at a notable cost to multiplicative capacity that abbreviates infections. Our results suggest that malaria parasites can enhance their fitness (cumulative host infectiousness) and persist longer within the host by modulating transmission investment over the course of infection. Organisms are typically expected to benefit from delaying reproductive investment early in life—to allow for increases in biomass—and terminally investing at the end of life. Our model indicates that human malaria parasites can likewise increase their fitness by delaying transmission investment initially and increasing investment at the end of infection. Thus, reproductive-survival tradeoffs can provide a general explanation for why pathogenic organisms do not evolve ever faster multiplication rates.

## 2 Materials and methods

We modified a previous model of human malaria infections Greischar et al. (2019) to incorporate realistic immune clearance and identify the resulting tradeoffs. As before, the model links transmission investment to its consequences for parasite fitness (*f*), i.e., cumulative host infectiousness to mosquitoes:

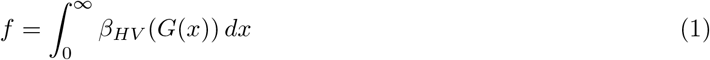

where *G* is the abundance of transmission stages (gametocytes) per microliter of blood at infection age *x*, and the proportion of mosquitoes infected (i.e., host infectiousness, *β*_*HV*_) is an empirically-derived curve fit to human infection data by Huijben et al. (2010):

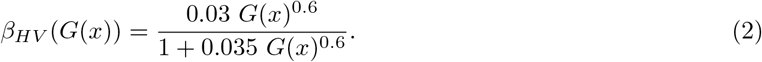

This quantity was recently referred to as the human-to-mosquito transmission probability Stopard et al. (2021) and the transmission efficiency in classic *R*_0_ formulations Smith and McKenzie (2004), but we use the term host infectiousness to emphasize the dependence on within-host gametocyte dynamics and the age of infections. Gametocyte dynamics over infection ages are specified by a coupled system of ordinary differential equations (ODEs) and partial differential equations (PDEs) as shown in Fig. 2A. Aside from infection age *x*, malaria biology encompasses two additional developmental delays that limit parasite capacity to multiply and transmit. First, following invasion of red blood cells (RBCs, *B*), parasites not allocated to gametocyte production must replicate within infected RBCs (iRBCs, *I*) for a period of time that is typically a multiple of 24 hours—48 hours for *P. falciparum* Mideo et al. (2013)—before iRBCs can burst to release RBC-invasive forms called merozoites (*M*) capable of initiating another round of multiplication in the blood. (For clarity, we use the term iRBC exclusively to refer to invaded RBCs allocated to within-host multiplication.) Second, some fraction of invaded RBCs (the transmission investment, *c*) are invested into the production of transmissible gametocytes (*G*) with the potential to infect blood-feeding mosquitoes and, subsequently, new hosts. Following invasion of an RBC, developing gametocytes (*I*_*G*_) require a lengthy period before maturing to become infectious, approximately 11 days for *P. falciparum* Lensen et al. (1999). *We therefore track the time since invasion of RBCs as the invasion age, τ* days (Table 1).

**Table 1:** Summary of model variables.

|  |  |
| --- | --- |
| $x$ | Infection age: time since start of blood-stage infection (days) |
| $\tau$ | Invasion age: time since invasion of RBCs (days) |
| $B(x)$ | Uninfected RBCs at infection age $x$ |
| $M(x)$ | Merozoites at infection age $x$ |
| $I(x, \tau)$ | Asexually infected RBCs (iRBCs) at infection age $x$ and invasion age $\tau$ |
| $I_G(x, \tau)$ | Developing gametocytes at infection age $x$ and invasion age $\tau$ |
| $G(x)$ | Mature gametocytes at infection age $x$ |
| $A(x)$ | Immune effectors at infection age $x$ |

**Figure 2:**
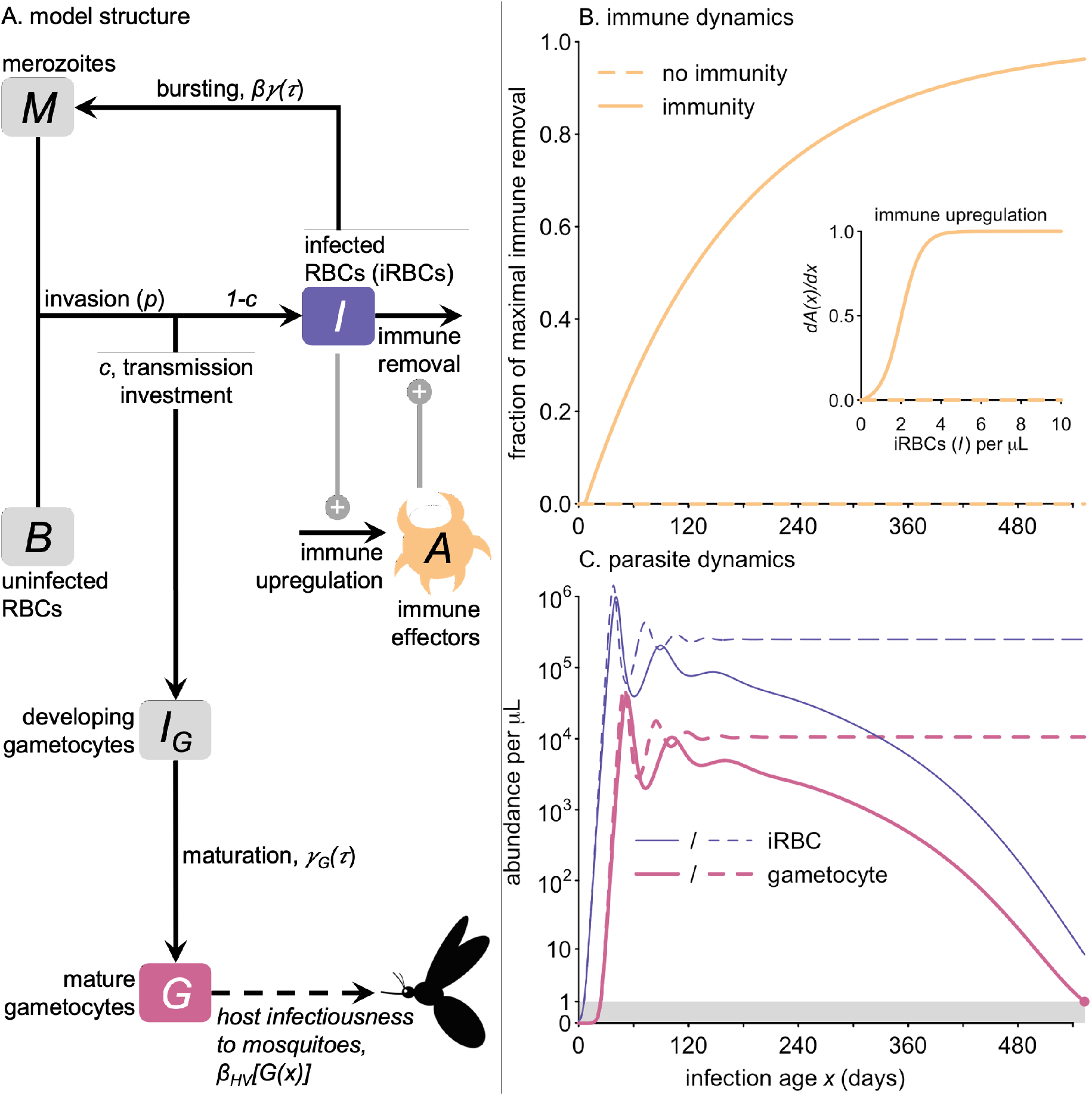
The model incorporates dynamic feedback between transmission investment (*c*) and immune clearance. (A) Within the host, merozoites (*M*) contact uninfected RBCs (*B*) and invade at rate *p*. Of the invaded RBCs, a fraction 1 − *c* becomes infected RBCs (iRBCs, *I*) that subsequently burst with rate *γ*(*τ*) where *τ* is the time since RBC invasion. When iRBC abundance exceeds a threshold, immune effector abundance (*A*) increases, increasing the removal rate of iRBCs. Surviving iRBCs burst to release *β* merozoites that can continue the proliferative cycle. A fraction *c* of invaded RBCs become developing gametocytes (*I*_*G*_) that mature into infectious gametocytes (*G*) at rate *γ*_*G*_(*τ*). Infectiousness, the probability of infecting mosquitoes, is an empirically derived function of gametocyte abundance (*β*_*HV*_ [*G*(*x*)]) at a given age of infection (*x*). (B) The rate of immune removal increases throughout infection (solid orange curves) as a function of iRBC (*I*) abundance (inset) or can be set to zero (dashed orange lines) to enable comparison with a previous version of this model Greischar et al. (2019). (C) Whatever the transmission investment (here *c* = 0.05), iRBCs (thin blue) and gametocytes (thick pink) undergo damped oscillations either to a nonzero equilibrium in the absence of immunity (dashed) or to zero with immunity (solid). We assume the infection ends (closed pink point) when gametocyte abundance declines to the minimum threshold thought to allow for transmission to mosquitoes (gray rectangle).

Following Greischar et al. (2019), we assume that uninfected RBCs (*B*(*x*)), are produced at a maximum rate denoted by *λ* with carrying capacity *K*, and removed at a background rate *µ* to maintain a homeostatic equilibrium in the absence of infection:

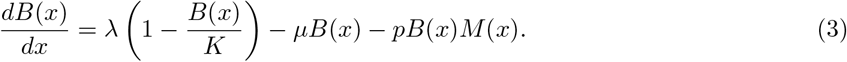

These uninfected RBCs are invaded by merozoites (*M* (*x*)), at invasion rate *p*. If not allocated to gametocyte production, invaded RBCs become iRBCs (*I*(*x, τ*)):

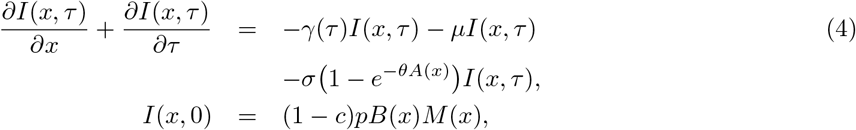

where *γ*(*τ*) is the rate at which iRBCs burst to release merozoites and *µ* is the background mortality rate for all RBCs (regardless of infection status). The rate of bursting (*γ*(*τ*)) is assumed to be gamma-distributed with a mean of two days (the time required for invasion to bursting for *P. falciparum*, Mideo et al. (2013)) and variance of 0.5 days. We extend the previous model formulation by adding a term for immune killing of iRBCs, where *σ* is the maximum rate at which iRBCs are killed by immune effectors (*A*(*x*)) modulated by a scaling parameter *θ* (details below in the section *Formulation of immunity*). If not removed by background or immune-mediated mortality, iRBCs burst to release merozoites:

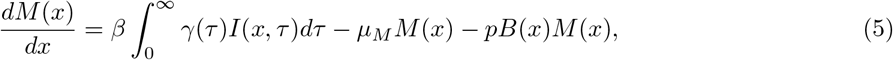

where *β* is the number of merozoites produced per iRBC. The background mortality rate of merozoites (*µ*_*M*_) is substantially higher than that of RBCs since merozoites are viable for only minutes after bursting Boyle et al. (2010).

A fraction *c* of invaded RBCs invested into transmission stage production become developing gametocytes (*I*_*G*_(*x, τ*)):

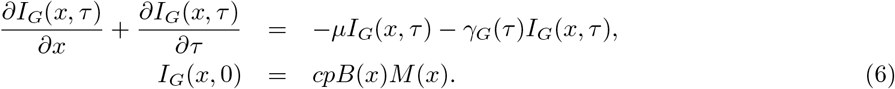

Developing gametocytes are assumed to be subject to the same background mortality rate of all RBCs (*µ*). However, immune killing is assumed to be negligible since most immune responses identified to date target iRBCs rather than developing or circulating gametocytes Gonzales et al. (2020); Bousema and Drakeley (2011). Developing gametocytes mature into infectious gametocytes (*G*(*x*)) at rate *γ*_*G*_(*τ*):

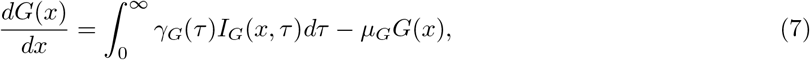

As with iRBC bursting, *γ*_*G*_(*τ*) is assumed to be gamma distributed with a mean of 11 days, the time required for developing gametocytes to reach full infectiousness *in vitro* Lensen et al. (1999), *and a variance of 3*.*67 days. Upon maturation, gametocytes are capable of infecting mosquitoes (Eq. 2)*. *We define the optimal transmission investment strategy c* as maximizing Eq. 1. We assume constant investment across all ages of infection and later relax that assumption to identify the best strategy when *c* varies with infection age *x*. Following previous work Greischar et al. (2016a), we specifying age-varying transmission investment *c* as a cubic spline with an intercept. We then compare those results with splines that are less flexible (linear, quadratic) or more flexible (with an interior knot) to test the impact of spline flexibility on fitness.

### 2.1 Formulation of immunity

Human malaria infections can persist for hundreds of days, exhibiting repeated waves of increase and decline in the iRBC abundance. The first peak in iRBC abundance is typically the largest with peak abundance declining in each subsequent wave until iRBCs are no longer detectable Eichner et al. (2001); Childs and Buckee (2015). Killing of iRBCs can occur via innate or general adaptive immunity that acts against all iRBCs, or by immune effectors that react (or cross-react) with specific antigens on the surface of iRBCs, which change over the course of infection as a parasite genotype cycles through its repertoire of antigenic variants Dzikowski et al. (2006); Recker et al. (2011); Noble et al. (2013); Day et al. (2017). Despite this complexity, Childs and Buckee (2015) determined that the pattern of declining peaks in iRBC abundance can only be recovered by a model that includes general adaptive immunity, which is increased when iRBC abundance exceeds a threshold, in effect tracking the cumulative exposure by the host to any parasite antigen. We adapt the formulation developed by Childs and Buckee (2015) to our continuous-time model framework by defining the increase of adaptive immunity as:

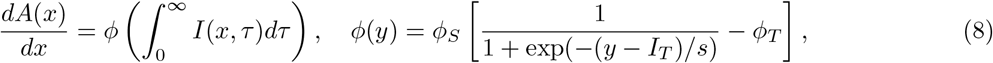

where *I*_*T*_ represents the threshold abundance of iRBCs (of any invasion age) above which immunity is increased. Eq. 8 results in scaling up of adaptive immune removal over the course of infection as immune upregulation increases in a sigmoidal fashion with increasing iRBC abundance (Fig. 2B and inset). We choose the constants *ϕ*_*S*_ and *ϕ*_*T*_ so that *ϕ*(0) = 0 and lim_*y*→∞_ *ϕ*(*y*) = 1. The previous model framework resulted in damped oscillations towards a nonzero equilibrium abundance of iRBCs and gametocytes within the host Greischar et al. (2019), but the addition of adaptive immune removal allows for reduction of iRBCs towards zero as infection age increases (Fig. 2C).

### 2.2 Infection length

We assume that infectiousness ends when gametocyte abundance declines to one per *µ*L. Since malaria parasites are obligately sexual, a mosquito blood meal must contain at least two gametocytes (one female and one male) to achieve fertilization and successfully colonize the vector. Sinden et al. (2007) assume a blood meal size of 2.13*µ*L, which translates to a lower limit of infectiousness of approximately one gametocyte per *µ*L. Thus, any relationship between transmission investment *c* and infection duration will emerge from model dynamics rather than being explicitly assumed in the model formulation. We compare the optimal strategies we find in our extended model framework with those found previously by either assuming a constant duration of infection *a*_*c*_ or enforcing an end to host infectiousness through the use of a constant recovery rate *ψ* until a maximum infection length *a*_*c*_ Greischar et al. (2019):

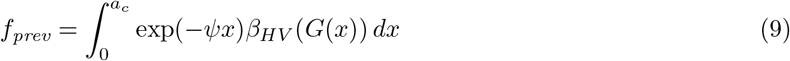

### 2.3 Merozoite effective reproductive number

We calculate an effective merozoite reproductive number, *R*_*M*_ (*t*) Pak et al. (2024), to clarify the role of immunity in limiting parasite population growth within the host, given as

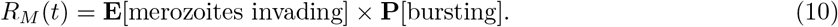

The expected number of merozoites invading is given by

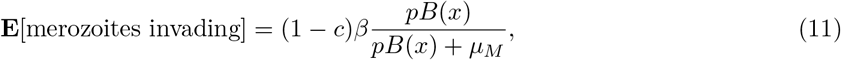

where the fraction indicates the probability that merozoites invade uninfected RBCs (*B*, which are invaded at rate *p*), as opposed to succumbing to background mortality, *µ*_*M*_ Boyle et al. (2010). Following invasion, iRBCs have some probability of surviving to burst successfully:

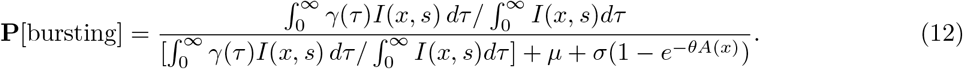

Since the rate of bursting depends on the invasion age *τ*, the numerator (and first term in the denominator) represents the weighted average rate of bursting (weighted by the invasion ages of the iRBCs present at infection age *x*). The other two terms in the denominator represent the loss of iRBCs to background RBC mortality (*µ*) and removal by adaptive immunity.

### 2.4 Sensitivity analysis

We first confirm that our PDE reformulation of the delayed differential equation system in Greischar et al. (2019) recovers the same optimal level of (constant) transmission investment under the same assumptions of either fixed infection duration or constant recovery rate *ψ*. We then compare optimal transmission investment strategies that are either constant across or vary with the age of infection *x*. To determine whether the shape of the optimal time-varying strategy depends on our choice of cubic spline, we re-ran optimizations assuming either lower or higher degrees of freedom (i.e., linear, quadratic, quartic). We also assess sensitivity to immune parameters and the initial inoculum size of iRBCs.

Previous theory suggests that rodent malaria parasites (*P. chabaudi*) should benefit from an initial delay before investing into transmission so that parasites can grow their numbers and subsequently produce gametocytes in much larger numbers, analogous to delayed reproductive investment in macroorganisms. Unlike *P. chabaudi*, human *P. falciparum* parasites produce substantially greater numbers of merozoites per iRBC (burst size), so we vary the burst size *β* and identify optimal constant and time-varying strategies.

### 2.5 Initial conditions & model simulation

We assume that blood-stage infection begins when iRBCs are inoculated into a host, mimicking the first cohort of iRBCs that are invaded following the latent liver stage of infection (as in Greischar et al. (2014, 2016a, 2019). For computational speed, we assume the ages (*τ*) of that initial inoculum (*I*_*init*_(0)) is uniformly distributed across invasion age *τ* from 0 to 48 hours, meaning an asynchronous start to infection. Past work suggested the level of synchrony in the initial inoculum (uniform versus narrow symmetric beta distributions) does not alter the optimal transmission investment strategy Greischar et al. (2016a). Hosts are assumed to begin infection with their homeostatic equilibrium RBC count *B*^*^ (Table 2). We assume that all hosts begin infection (*x* = 0) with no merozoites, gametocytes (developing or mature), or adaptive immune effectors (*M* (0) = *I*_*G*_(0) = *G*(0) = *A*(0) = 0). The numerical scheme for simulating the model framework is described in Appendix A. We performed a convergence study to verify the accuracy of the method and used a step size of *h* = 0.125 for all results.

**Table 2:** Within-host parameter values & units for human malaria infections.

| Parameter | Definition | Value | Source |
| --- | --- | --- | --- |
| $B^*$ | RBC count at infection-free equilibrium | $5 \times 10^6$ cells/μL | Loe and Edwards (2004) |
| $K$ | carrying capacity of RBCs | $\lambda B^*/(\lambda + \mu B^*)$ cells/μL | Determined by $B^*$ |
| $\lambda$ | max. new RBCs | $2 \times 10^5$ RBCs/μL/day | Koury and Ponka (2004) |
| $\mu$ | RBC mortality rate | 1/120 /day | Koury and Ponka (2004) |
| $p$ | per merozoite invasion rate | $8.35 \times 10^{-6}$/day | Unknown for *P. falciparum* parasites, within the range reported for *P. chabaudi* Mideo et al. (2008) |
| $\beta$ | burst size | 16 merozoites | Reilly et al. (2007) |
| $\mu_M$ | merozoite mortality rate | 200/day | Boyle et al. (2010) |
| $\mu_G$ | mortality rate of mature gametocytes | 0.5/day | maximally infectious period of two days Lensen et al. (1999) |
| $\sigma$ | maximum rate of immune removal of iRBCs | 0.55/day | — |
| $I_T$ | iRBC abundance threshold that triggers immune upregulation | 2 iRBCs/μL | Childs and Buckee (2015) estimate threshold as $10^7$ iRBCs/human |
| $\theta$ | scaling of immunity level | $2.5 \times 10^{-4}$/immune effector $A$ | — |
| $s$ | immune sensitivity to iRBC abundance | 1 | — |
| $I_{init}(0)$ | abundance of iRBCs at infection age $x = 0$ | $\int_0^\infty I(0, \tau) d\tau = 0.06/\mu$L | Assuming mosquito injects 10 sporozoites, each infecting a hepatocyte and releasing $3 \times 10^4$ merozoites that generate iRBCs Paul et al. (2003) |

## 3 Results

### 3.1 Transmission investment reduces infection duration

With the inclusion of plausible immune clearance, infections can end via immune removal and both the degree and duration of infectiousness vary with transmission investment (Fig. 3). Increasing transmission investment shortens simulated infections when we incorporate general adaptive immune clearance inspired by Childs and Buckee (2015) (Fig. 3A). As before Greischar et al. (2019), we find that lower transmission investment hastens the peak in infectiousness by enabling more rapid expansion of iRBC numbers within the host. In the absence of immune clearance (*σ* = 0), we recover the same optimal levels of transmission investment found in the previous delayed differential equation formulation Greischar et al. (2019). Specifically, for maximum infection duration of *a*_*c*_ = 280 days with no host recovery, we find that a transmission investment *c* ∼ 20% maximizes *f*_*prev*_ (Eq. 9). Assuming a constant rate of recovery *ψ* = 1*/*105 (independent of any within-host dynamics), we recover the result that transmission investment *c* ∼ 15% maximizes fitness (*f*_*prev*_). In contrast, a much lower level of transmission investment (*c* ∼ 4%) is favored when we incorporate immune effectors so that host recovery rates represent the outcome of within-host dynamics (Fig. 3B). We find that restrained investment into transmission prolongs infection duration (Fig. 3C), consistent with the observation from human *P. falciparum* cases that infections tend to persist longer when parasites exhibit faster multiplication rates Mackinnon and Read (2004). Thus, we predict that immunity imposes a survival-reproduction tradeoff on malaria parasites such that greater transmission investment reduces persistence within the host.

**Figure 3:**
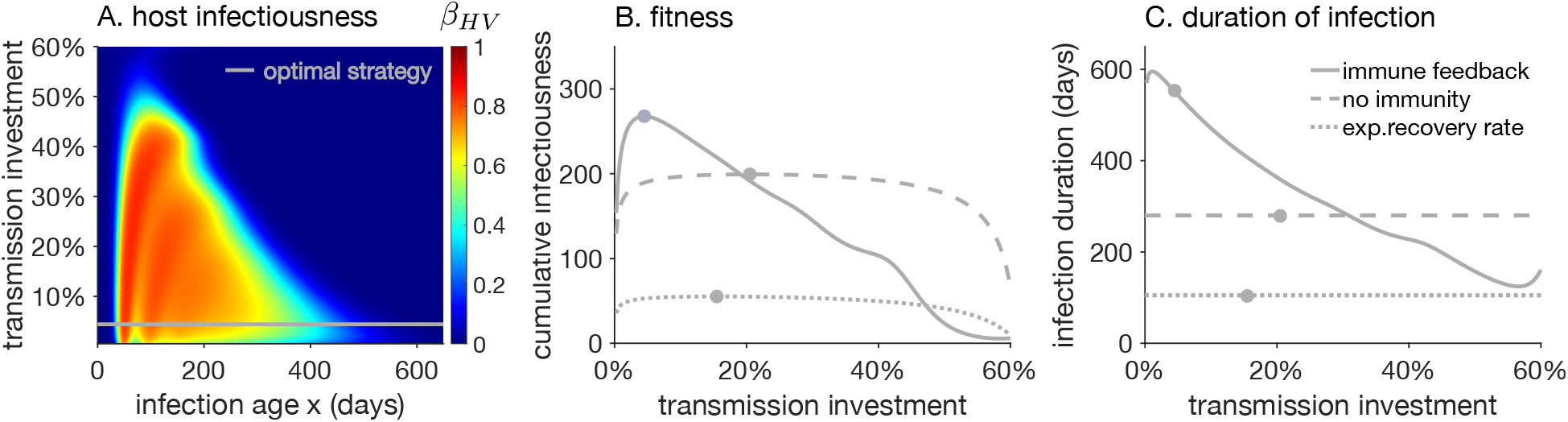
With the addition of plausible immune clearance of parasites, greater investment in transmission reduces the duration of host infectiousness. (A) The probability of transmitting an infection to mosquitoes (*β*_*HV*_) is shown as a heatmap over the days since infection (x-axis) for varying transmission investment (y-axis), showing that lower transmission investment levels extend the time the host is highly infectious. The optimal constant investment strategy for maximizing the probability of onward transmission (indicated by a white horizontal line) is *c* = 4.4%. (B) Cumulative infectiousness (*f*, the infection probability integrated over the duration of infection) is shown across increasing levels of constant transmission investment. The solid line indicates the full model with the potential for the level of transmission investment to alter the time until immune clearance. The other curves show cumulative infectiousness for the same model without immunity (i.e., *σ* = 0) calculated according to Eq. 9 with the maximum infection length set to *a*_*c*_ = 280 days Greischar et al. (2019), either assuming no recovery (dashed, *ψ* = 0) or a constant rate of recovery (dotted, *ψ* = 1*/*105 days). The infection lengths corresponding to each scenario are shown in (C). Closed dots indicate the optimal level of transmission investment for each set of model assumptions.

We find that increasing levels of transmission investment alter effective merozoite numbers (*R*_*M*_) and by extension, cumulative infectiousness (fitness, *f*) and infection duration in two distinct ways. First, transmission investment reduces the capacity for within-host multiplication, as seen from the expected merozoites invading (Fig. 4A and in line with past theory Mideo et al. (2008); Koella and Antia (1995); Greischar et al. (2016a, 2019). Second, by reducing the number of iRBCs produced, transmission investment reduces immune upregulation (Fig. 1B, inset) and thereby increases the probability of successfully bursting despite immune pressure (Fig. 4B). However, the negative impact on multiplicative capacity is noticeable even with small increases in transmission investment, while the reduction in immune removal is barely discernible, so that the net effect of transmission investment is to reduce infectiousness and abbreviate infections. As a result of these competing impacts, the duration of infection initially increases when transmission increases up to 1%, which yields the maximum duration of infection (but not the maximum fitness). Increasing transmission investment further yields a rapid decrease in infection duration (Fig. 3C).

**Figure 4:**
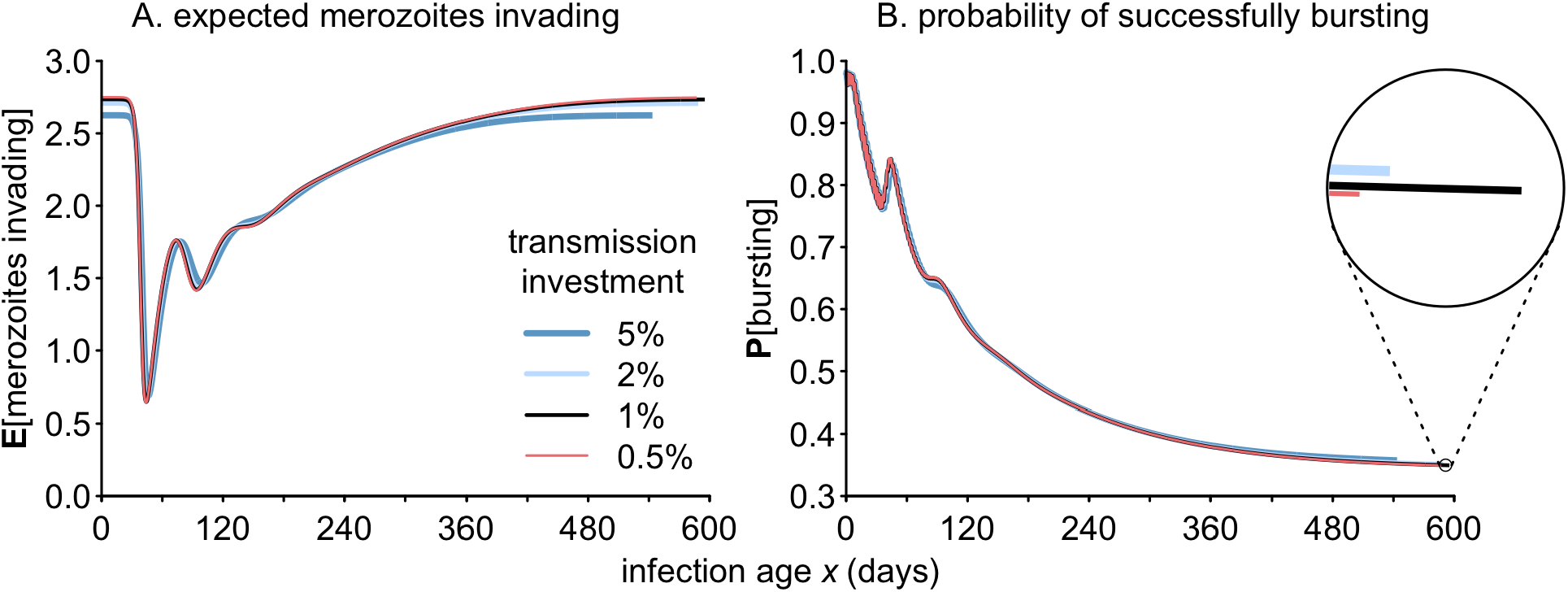
Transmission investment reduces capacity for within-host multiplication. Modest increases in the constant level of transmission investment cause (A) noticeable reductions in the expected merozoites invading, and (B) very small reductions in the rate of removal by immunity. Note that 1% transmission investment (black) yields the longest infection for our parameterization, but not the greatest fitness.

### 3.2 Age-varying strategies outperform constant transmission investment

We find that transmission investment strategies that vary with the age of infection (*x*) generate longer simulated infections with greater cumulative infectiousness (*f*, Fig. 5). Lower initial transmission investment hastens the increase in infectiousness, in line with theory for *P. chabaudi* in the absence of adaptive immunity Greischar et al. (2016a). A smaller fraction of iRBCs allocated to transmission can produce a larger number of transmissible gametocytes due to increased multiplicative capacity (Fig. 4A). Hence the cubic spline strategy with a knot exhibits a faster accumulation of infectiousness compared with the constant strategy (Fig. 5B).

**Figure 5:**
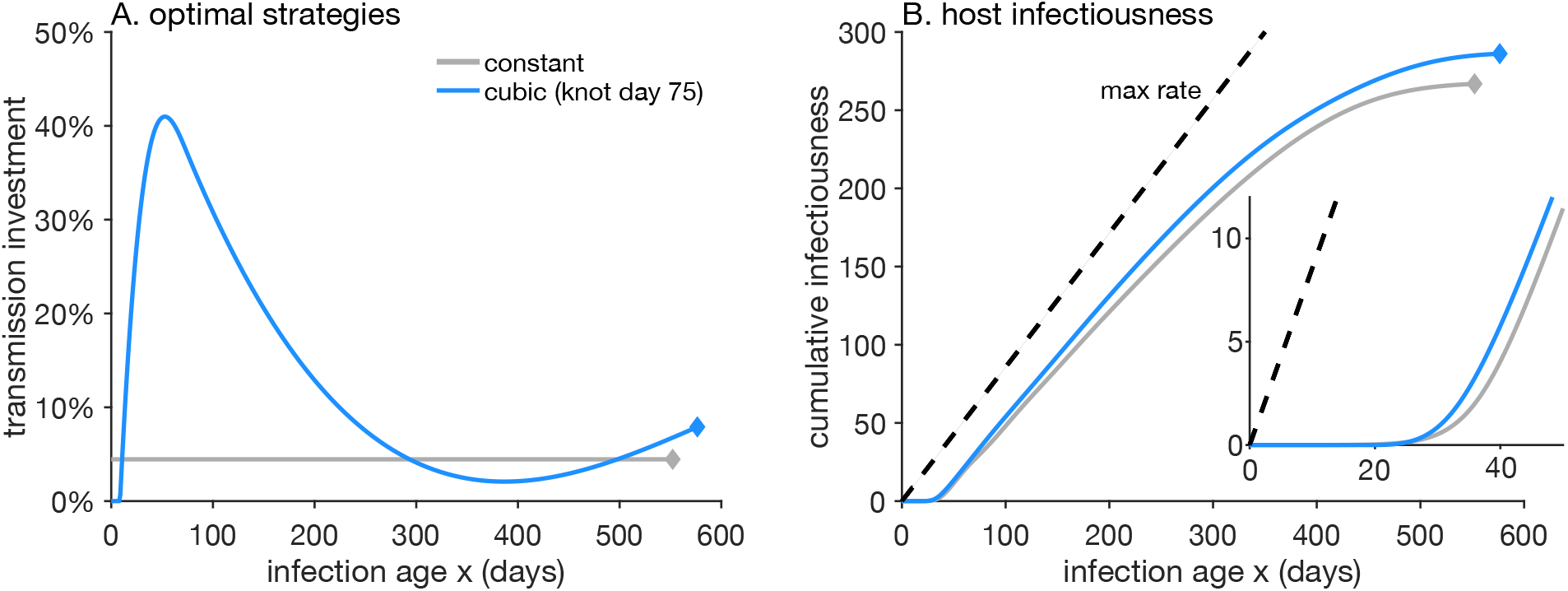
Time-varying transmission investment strategies outperform constant ones. (A) Transmission investment is initially delayed for sufficiently flexible splines (blue, cubic spline with knot at day 75). The optimal constant strategy is shown in gray for comparison. (B) Time varying strategies yield higher fitness returns over the full infection duration. Cumulative host infectiousness increases most rapidly when initial transmission investment is delayed (blue, compare with constant investment in gray). Closed diamonds indicate the point the infection drops below one gametocyte per *µ*L, the assumed threshold for transmission to mosquitoes.

### 3.3 Sensitivity to burst sizes, initial inoculum & spline flexibility

We find that increasing burst size (*β*) enhances the potential fitness gains and extends infection duration (Fig. S1). Even small reductions in burst sizes (e.g., from 16 to 14 or 12) can shorten infections by a hundred or more days and dramatically reduce cumulative infectiousness. Increasing the initial inoculum does not alter the optimal fixed transmission investment strategy, though larger initial abundance increases the maximum cumulative infectionsness (Fig. S2). Similarly, the initial inoculum size does not qualitatively change the optimal age-varying strategy but does allow higher initial levels of transmission investment in the optimal strategy (Fig. S3).

We find that cumulative infectiousness increases but saturates with spline complexity (Fig. S4), consistent with a previous study using splines to describe optimal transmission investment strategies in *P. chabaudi* Greischar et al. (2016a). Adding an interior knot to the spline increases flexibility and hence cumulative infectiousness over that of the optimal cubic strategy without a knot (Fig. S5. Including a knot modestly increased infection duration by 5-6 days depending on the knot placement (Fig. S5B). The placement of the knot causes minor differences in cumulative infectiousness, with a knot at day 75 showing a narrow advantage compared to knots at day 25 or day 125. All splines with sufficient flexibility (i.e., for infections lasting more than 500 days, with a knot) exhibit an initial delay in transmission investment (Fig. S5). Knot placement influences the duration of the initial delay in transmission investment, with earlier knots allowing for longer delays (Fig. S5A). Due to the reduction in infection duration, the spline flexibility over the duration of infection increases as burst size declines. Transmission investment strategies specified as a cubic spline with no knot allowed greater flexibility over shorter versus longer infections so that lower burst sizes also demonstrate initial delays in transmission investment that are predicted to be optimal. As a result of that delayed transmission investment, lower burst sizes exhibit a faster increase in infectiousness compared with the optimal cubic strategy without a knot for the baseline burst size *β* = 16 (Fig. S1).

## 4 Discussion

Tradeoffs between the rate and duration of transmission form the basis for most theory regarding the evolution of pathogenic organisms (Anderson and May (1982); Frank (1996); Gandon et al. (2001); Alizon (2008); Saad-Roy et al. (2020, 2021a,b); Miller and Metcalf (2022); Ripoll and Font (2023), reviewed in Bull and Lauring (2014); Alizon and Sofonea (2021)). In the absence of such tradeoffs, it is not obvious what constrains the fitness of pathogenic organisms, nor what would prevent the evolution of ever faster rates of within-host multiplication. We find that incorporating adaptive immunity into a within-host model of human malaria infections constrains parasite fitness by imposing a reproduction-survival tradeoff. Thus, we show that transmission-duration tradeoffs are not required to constrain parasite fitness. Instead, our results suggest that reproduction-survival tradeoffs may represent a more general constraint on the evolution of both pathogenic and free-living organisms.

Our model framework reconciles two seemingly contradictory patterns in malaria biology. First, faster multiplication rates have been observed to extend the time until immune clearance Mackinnon and Read (2004) but slower multiplication and lower iRBC abundance is typically thought to reduce pressure from the immune system Borrmann and Matuschewski (2011); Mideo et al. (2011); Andrade et al. (2020). Our model shows that both of these assumptions can be true: reducing transmission investment boosts within-host multiplication rates and thereby extends the lifespan of infection, even while it reduces the probability of surviving immune clearance. The key to reconciling these patterns is that transmission investment-mediated reductions in multiplication rates dramatically alter within-host growth potential, while inducing extremely minor reductions in the probability of surviving immune clearance. Our analysis suggests that other components of multiplicative capacity—especially increasing burst sizes—may likewise extend the duration of infection even as they hasten upregulation of host immunity. The second apparent and long-standing contradiction is that both human and rodent malaria parasites appear to invest far less into transmission than would seem to be optimal Taylor and Read (1997). Within-host models consistently find optimal levels of transmission investment far in excess of values inferred from data (typically less than 10%, Eichner et al. (2001), Greischar et al. (2016b)), unless they invoke competition from a coinfecting strain McKenzie and Bossert (1998); Mideo and Day (2008); Greischar et al. (2016a, 2019); Pak et al. (2024) or evolutionary feedbacks from epidemiological dynamics Greischar et al. (2019). We show that incorporating a plausible formulation of adaptive immunity can select for optimal levels of transmission investment much closer to those deemed reasonable, even in the absence of coinfection or selection from multiple scales. Thus, the dynamics of adaptive immunity represent a key source of selection on transmission investment strategies.

Motivated by the relatively low fraction of malaria cases that end due to infection-induced death of the host Cunnington et al. (2013); Lindblade et al. (2013), our model assumes that clearance by the immune system is the primary way human malaria infections end. Infections can also end by curative drug treatment, and accounting for drug-induced clearance will be important to adapting this approach to low transmission settings. Infections may be more often drug treated in low transmission areas where less frequent exposure means that infected individuals more often exhibit acute symptoms. In areas with high infection prevalence where frequent exposure reduces symptoms Lindblade et al. (2013), malaria infections are less likely to be detected, and subsequently drug-treated. Our results suggest that parasites experience large fitness returns from the later portion of infections, but greater transmission investment makes it less likely that parasites will persist long enough to reap those fitness returns. Hence we find selection for reduced transmission investment. All else being equal, lower transmission investment should pose a greater risk to host health by enabling the greater within-host abundance associated with adverse outcomes Cunnington et al. (2013). Frequent drug treatment could negate those late-infection fitness returns and weaken selection for reduced transmission investment. However, robust predictions require information regarding a current unknown, how (or whether) transmission investment—by altering multiplication rates—modulates the onset of symptoms and by extension the timing of drug treatment. A better understanding of the within-host causes of symptom onset would enable models to explore plausible scenarios for evolution in response to widespread drug treatment in areas moving towards elimination.

By reducing both infection duration and the prevalence of coinfections, drug treatment has been predicted to select for increased transmission investment, with concerns that could enhance transmission potential and undermine the health gains of widespread drug treatment Early et al. (2022). In contrast, our results show strongly that increased transmission investment reduces transmission potential (i.e., cumulative infectiousness) and infection duration. We find that shortened infections are likely to select for reduced transmission investment, at least initially, since that hastens early increases in host infectiousness, a result that holds even in the absence of dynamic feedbacks with immunity and age-varying transmission investment Greischar et al. (2019). There is also reason to believe the drug treatment could have simultaneous distinct and conflicting impacts on infection duration. Widespread drug treatment also reduces the prevalence of coinfections by limiting transmission Greischar et al. (2020), thereby removing a potent selection pressure for reduced transmission investment McKenzie and Bossert (1998); Mideo and Day (2008); Greischar et al. (2016a, 2019). Accordingly, genes initiating gametocyte production showed greater expression in low versus high transmission zones Rono et al. (2018). Similarly, increasing frequency of alleles initiating gametocyte production was associated with a long-term decline in transmission in French Guiana, corresponding to intensive intervention efforts Early et al. (2022). Widespread drug treatment could also—by reducing transmission intensity—remove another common route by which persistence within the host is ended, displacement by a newly-inoculated *P. falciparum* strain Childs and Buckee (2015). As a result of strain displacement, infections by a particular genotype may only last days or weeks in high transmission zones Daubersies et al. (1996). In low transmission zones where coinfections are rare, most infections end via drug treatment or natural clearance, and increasing rates of drug treatment is likely to shorten mean infection duration. In high transmission zones, increasing rates of drug treatment may reduce infection length of treated infections but extend the duration of untreated infections by reducing the frequency of strain displacement. Multiscale models are needed to disentangle the relative importance of these two processes and identify how drug treatment will impact the evolution of transmission investment.

Our analysis lends further support to the idea that age-varying transmission investment strategies can outperform constant ones. The specifics vary with epidemiological context, which determines how much infectiousness at different infection ages contributes to parasite fitness Day (2003); Greischar et al. (2019). Placing our model results in the context of past theory suggests that multiple epidemiological scenarios are likely to select for reduced transmission investment. Past work found that an expanding epidemic should select for decreased transmission investment, with an even sharper decrease predicted when rates of host recovery were assumed to increase with investment in transmission Greischar et al. (2019). In an expanding epidemic, early infection ages are overrepresented, so that parasite traits that enhance transmission early in infection are favored. Since early reductions or delays in transmission investment hasten host infectiousness, that strategy should be favored when either epidemiology or mass drug administration increase the importance of transmission from early infection ages Greischar et al. (2019). Seasonality can impose different constraints on the timing of transmission, especially when mosquitoes are absent during regular dry seasons. Recent investigation of asymptomatic human infections suggest parasites face substantial immune pressure during the dry season that makes it challenging to persist within hosts until the onset of the wet season Andrade et al. (2020, 2024). Enhanced infectiousness during the dry season is a dead end and withinhost persistence is be a major component of parasite fitness. Even in such seasonal environments, our results suggest that a lower (constant) level of transmission investment is favored, since that dramatically increases the ability of parasites to increase within-host abundance and persist despite immune pressure. The optimal age-varying transmission investment strategy for a seasonal environment is more difficult to predict, since *P. falciparum* clones transmitted early in the wet season are less likely to persist to the end of the dry season Andrade et al. (2024). Nonetheless, our results suggest that any traits that enhance multiplicative capacity should help parasites persist in spite of immune pressure.

While our model suggests parasites should benefit from changing transmission investment with infection age, the benefits of such flexible strategies vary depending on how quickly organisms can sense and respond to changing conditions Padilla and Adolph (1996). By setting transmission investment as a function of infection age, our model implicitly assumes that parasites have perfect information about the progression and end of infections. Thus, our predictions represent ideal strategies that may not be achievable given constraints on parasites’ capacity to sense and respond to their within-host environment. Even with the assumption of perfect information, we find that optimal strategies are sensitive to the flexibility of the spline per time unit, suggesting that much depends on how quickly parasites can alter their levels of transmission investment. Experimental infections suggest that the rodent malaria parasite *P. chabaudi* can alter transmission investment in response to its own within-host multiplication rate and availability of RBCs to infect Schneider et al. (2018), consistent with changes in transmission investment for *P. falciparum in vitro* Bruce et al. (1990). *Avian malaria parasites appear capable of responding to the cue of biting by uninfected mosquitoes Cornet* *et* *al*. *(2014)*. A more complete understanding of the cues that trigger changes in transmission investment—and how quickly—would help resolve the question of how closely parasites can match putative optimal strategies.

Tradeoffs between reproduction and survival can emerge as a result of pressure from natural enemies like pathogenic organisms (e.g., exposure to bacterial antigens in birds, Velando et al. (2006); frequent worm infections in Soay sheep, Graham et al. (2010)). Here we demonstrate that pathogenic organisms themselves can be subject to reproduction-survival tradeoffs due to pressure from the immune system, a prediction relevant to the diverse array of bacteria, fungi, and protozoa that produce specialized stages for within-host multiplication versus onward transmission Anderson and May (1981). For pathogenic organisms lacking such specialization, reproduction-survival tradeoffs are equivalent to the transmission-duration tradeoffs that have formed the basis for much evolutionary theory (May and Anderson (1983); Frank (1996); Gandon et al. (2001); Day (2003); Alizon (2008); Saad-Roy et al. (2020, 2021b,a); Miller and Metcalf (2022); Ripoll and Font (2023), reviewed in Bull and Lauring (2014); Bull and Antia (2022)). Thus, reproduction-survival tradeoffs offer a broadly relevant explanation for constrained evolution in both free-living and pathogenic organisms.

## A Appendix A

**Numerical Scheme**

The within-host dynamics involve two time scales: *x* and *τ* .

### Uninfected RBCs

The uninfected RBCs are described by:

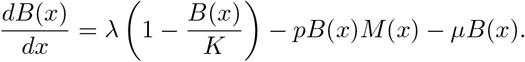

Assuming that *x* ∈ [0, *T*] ≈ {0, *h*, 2*h*, …, *Nh*}, *N* ∈ ℕ is divided by step *h* into intervals, discretizing for the *n*th interval gives

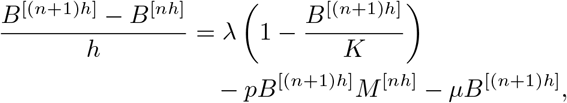

which rearranges to

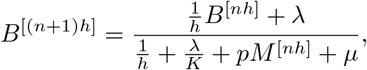

for *n* ≥ 0. The initial condition is *B*^[0]^ = *B*_0_ *>* 0.

#### Merozoites

The merozoites are described by:

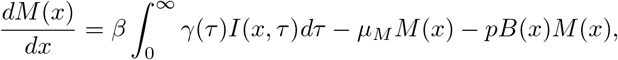

with *M* (0) = 0. Assuming that (*x, τ*) ∈ [0, *T*]^2^ ≈ {0, *h*, 2*h*, …, *Nh*}^2^, *N* ∈ ℕ is divided by step *h* into intervals, discretizing for the *n*th interval in *x* and the *j*th interval in *τ* gives

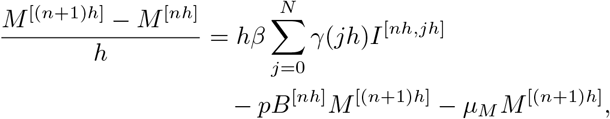

which rearranges to

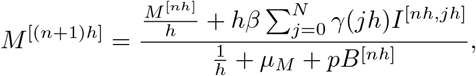

for *n* ≥ 0 and *j* ≥ 0. The initial condition is *M* ^[0]^ = 0. Note that we need *I*^[*nh*,*jh*]^∀*j* ∈ {1, …, *N*}.

#### Asexually infected RBCs

The asexually infected RBCs are described by

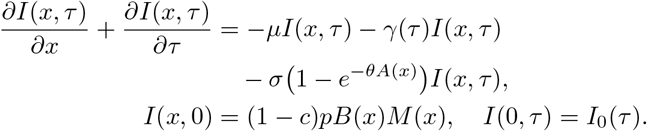

Discretizing for the *n*th interval in *x* and the *j*th interval in *τ* gives

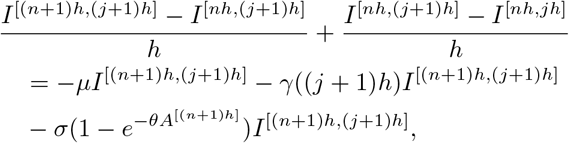

which rearranges to

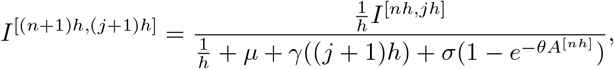

for *n* ≥ 0 and *j* ≥ 0. The initial conditions become *I*^[*nh*,0]^ = (1 − *c*)*pB*^[*nh*]^*M* ^[*nh*]^ for *n* ≥ 0 and *I*^[0,*jh*]^ = *I*(0) is a given distribution.

#### Immature sexually infected RBCs

The immature sexually infected RBCs are described by

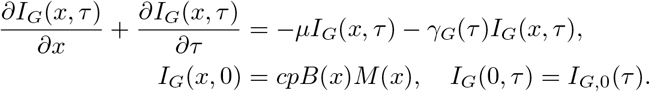

Discretizing for the *n*th interval in *x* and the *𝓁*th interval in *τ* gives

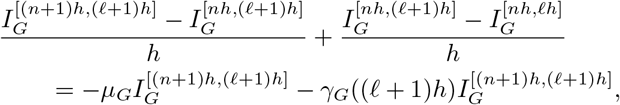

which rearranges to

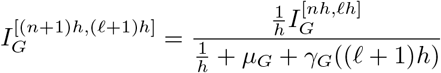

for *n* ≥ 0 and *𝓁* ≥ 0. The initial conditions become 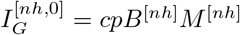 for *n* ≥ 0 and 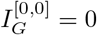.

#### Mature sexually infected RBCs

The mature sexually infected RBCs are described by

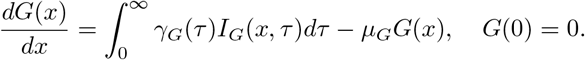

Discretizing for the *n*th interval in *x* and the *𝓁*th interval in *τ* gives

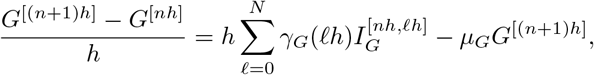

which rearranging to

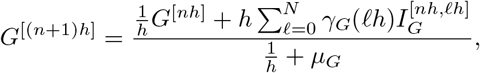

for *n* ≥ 0 and *𝓁* ≥ 0. The initial condition becomes *G*^[0]^ = 0. Note that we need 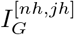 for *n* ≥ 0 and *j* ≥ 0.

#### Immunity levels

The immunity levels are described by

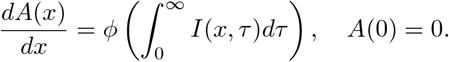

Discretizing for the *n*th interval in *x* and the *j*th interval in *τ* gives

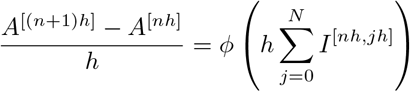

which rearranges to

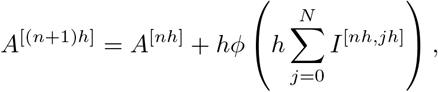

for *n* ≥ 0 and *j* ≥ 0. The initial condition becomes *A*^[0]^ = 0. Note that we need *I*^[*nh*,*jh*]^ for *n* ≥ 0 and *j* ≥ 0.

## B Supplemental figures

**Figure S1:**
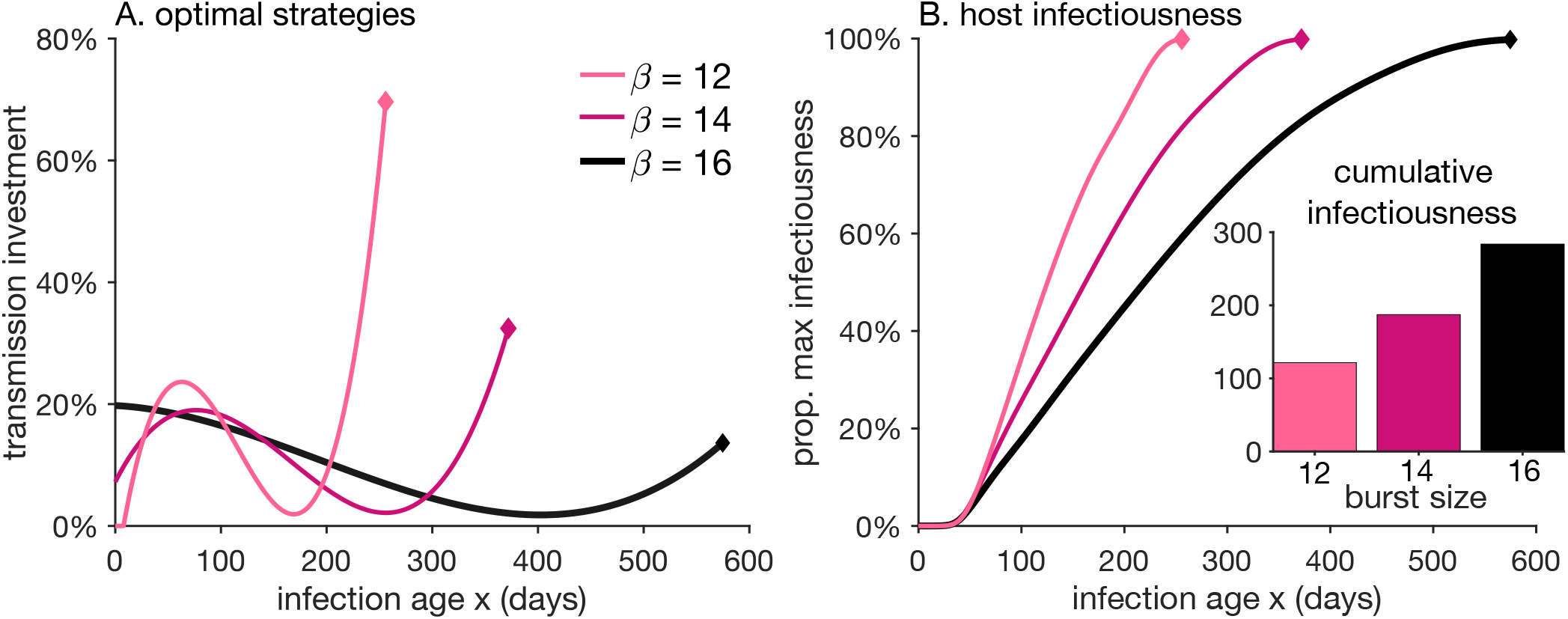
(A) Optimal time varying strategies with cubic splines (no knot) for different burst sizes (*β*). (B) Trajectories showing how quickly each strategy reaches its own maximum cumulative infectiousness over the course of infection; the inset shows the cumulative infectiousness value (*f*) for each strategy.

**Figure S2:**
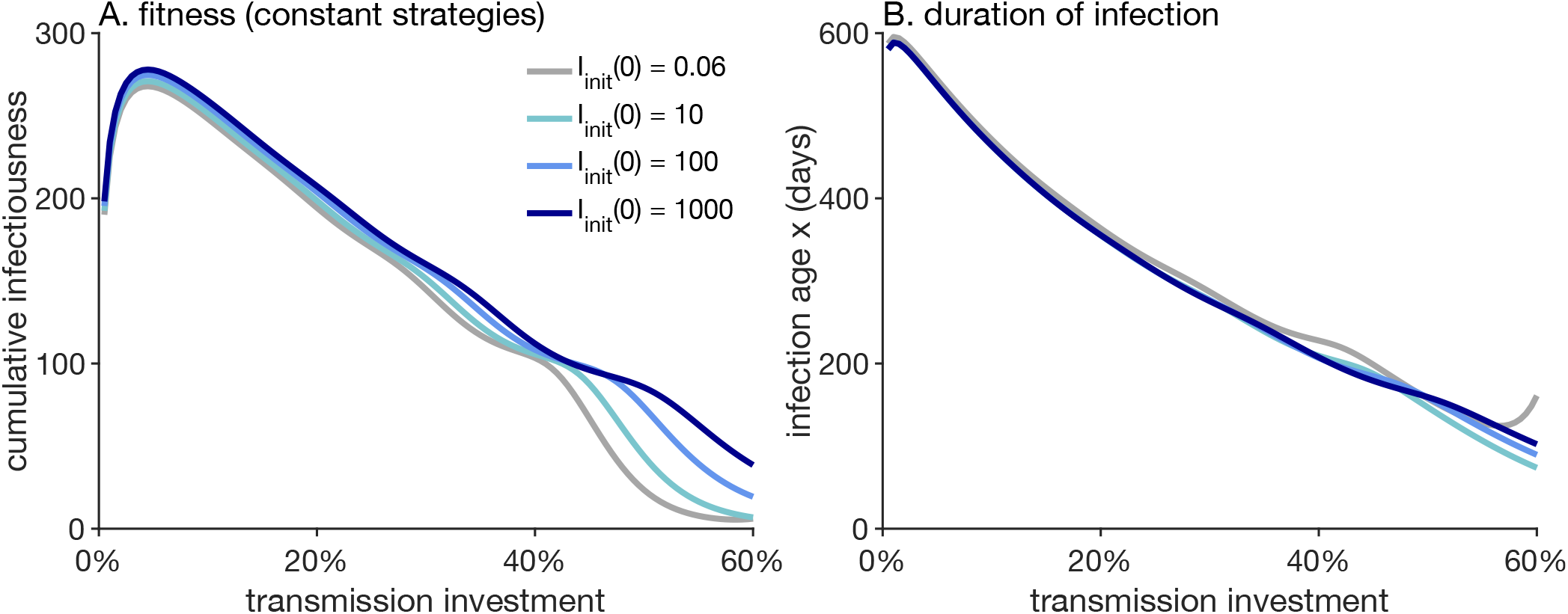
(A) Fitness profiles for fixed constant transmission investment strategies (i.e. values of the parameter *c*) for different values of the initial inoculum 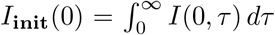. (B) Duration of infection versus transmission investment for different values of the initial inoculum.

**Figure S3:**
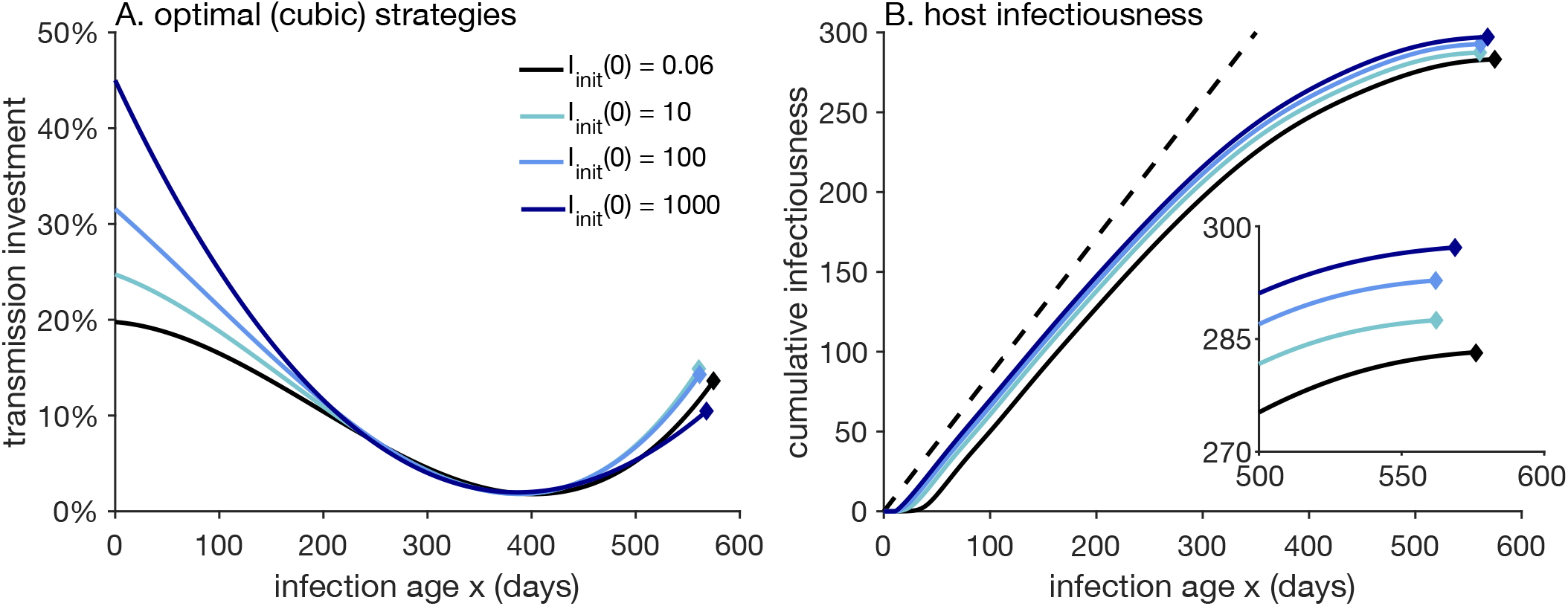
(A) Optimal strategies with cubic spline (no knot) for different values of the initial inoculum 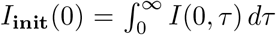. (B) Cumulative infectiousness (fitness) for different values of the initial inoculum as a function of age of infection (x).

**Figure S4:**
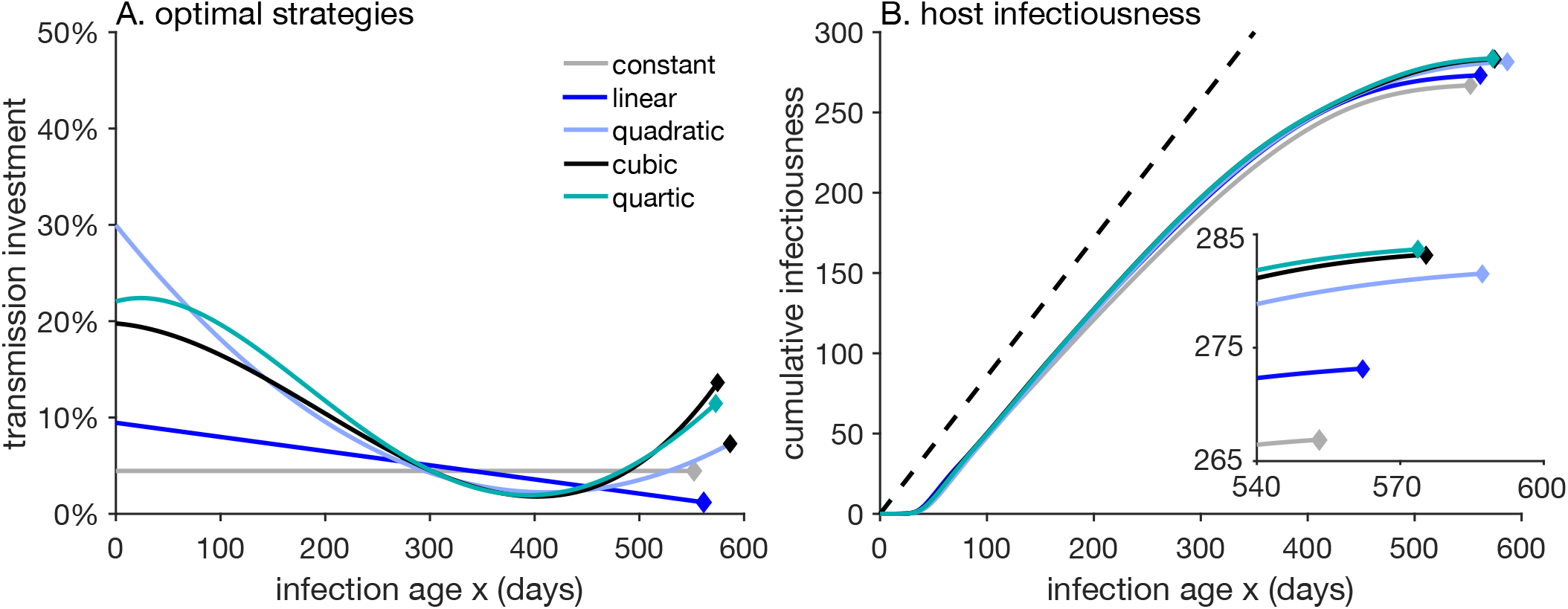
(A) Optimal time varying strategies with polynomial splines of different degrees. (B) Cumulative infectiousness versus time for the strategies shown in A.

**Figure S5:**
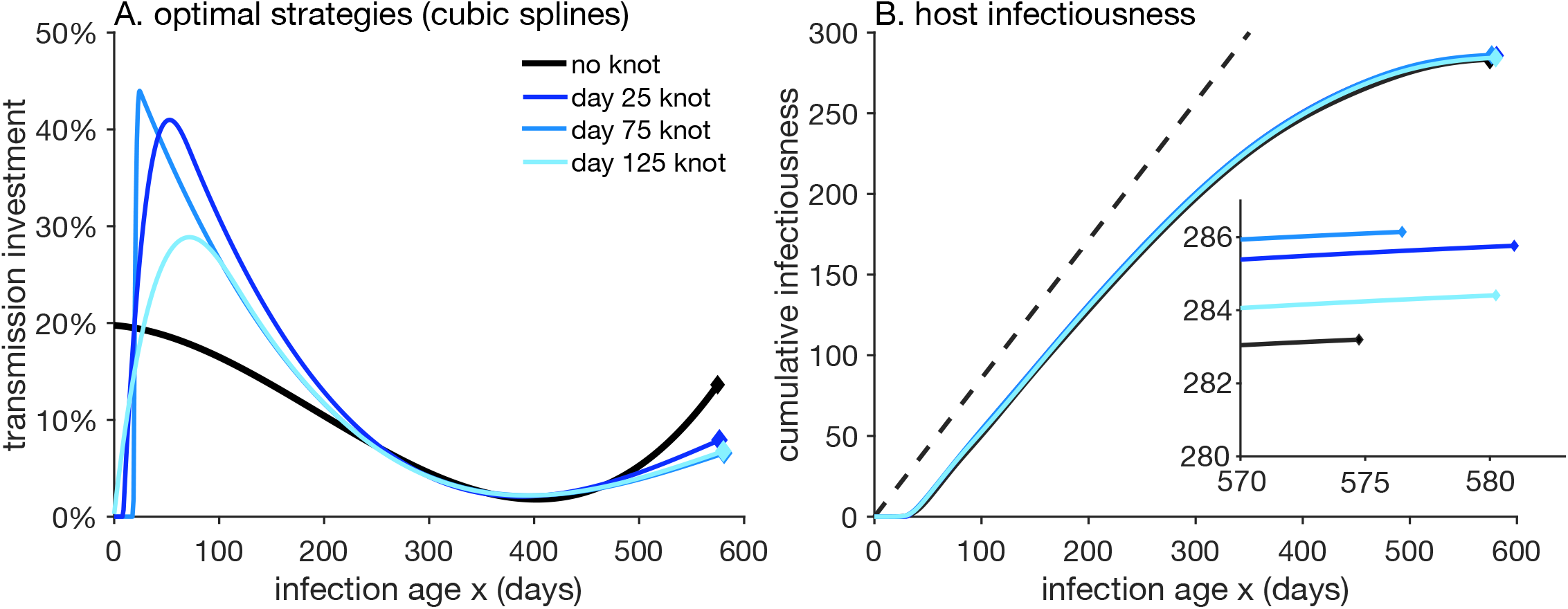
(A) Optimal strategies with cubic splines with different knot placements. (B) Cumulative infectiousness versus time for the strategies shown in A.

